# Histone H2AK119 Mono-Ubiquitination is Essential for Polycomb-Mediated Transcriptional Repression

**DOI:** 10.1101/690461

**Authors:** Simone Tamburri, Elisa Lavarone, Daniel Fernández-Pérez, Marika Zanotti, Daria Manganaro, Eric Conway, Diego Pasini

## Abstract

The major function of Polycomb group proteins (PcG) is to maintain transcriptional repression to preserve cellular identity. This is exerted by two distinct repressive complexes, PRC1 and PRC2, that modify histones by depositing H2AK119ub1 and H3K27me3, respectively. Both complexes are essential for development and are deregulated in several types of human tumors. PRC1 and PRC2 exist in different variants and show a complex regulatory cross-talk. However, the contribution that H2AK119ub1 plays in mediating PcG repressive functions remains largely controversial. Coupling an inducible system with the expression of a fully catalytic inactive RING1B mutant, we demonstrated that H2AK119ub1 deposition is essential to maintain PcG-target genes repressed in ESC. Loss of H2AK119ub1 induced a rapid displacement of PRC2 activity and a loss of H3K27me3 deposition. This affected both PRC2.1 and PRC2.2 variants and further correlated with a strong displacement and destabilization of canonical PRC1. Finally, we find that variant PRC1 forms can sense H2AK119ub1 deposition, which contributes to their stabilization specifically at sites where this modification is highly enriched. Overall our data place H2AK119ub1 deposition as central hub that mount PcG repressive machineries to preserve cell transcriptional identity.

## INTRODUCTION

Organism development and adult tissue homeostasis requires a precise and dynamic control of cellular transcriptional identity. Several chromatin remodelling activities contribute to the establishment of precise transcriptional states by modifying the chromatin environment. This also involves the regulation of post-translational modifications of histone proteins by highly specialized enzymes that by “writing”, “reading” and “erasing” specific modifications define the transcriptional state of target genes (Jenuwein and Allis, 2001; Pasini et al., 2008). Consistent with their essential role in controlling cellular identity, these mechanisms also play critical roles in the development of different human pathologies, with cancer being a leading example. Indeed, chromatin modifiers represent one of the most frequently mutated group of genes across all types of human tumours (Comet et al., 2016; Flavahan et al., 2017; Pasini and Di Croce, 2016).

Polycomb Group proteins (PcG) play a central role in these processes and represent the major repressive mechanism utilized in facultative heterochromatin (Bracken and Helin, 2009; Scelfo et al., 2015). PcGs were first discovered in *Drosophila melanogaster* where they play an essential role in maintaining the correct spatio-temporal repression of homeotic genes during fly development (Paro, 1990). This repressive function has been maintained in mammals where PcGs contribute to the repression of all CpG island (CpGi) containing promoters (Mendenhall et al., 2010; Riising et al., 2014). This involves the cooperative activity of two large Polycomb repressive complexes termed PRC1 and PRC2. Both complexes are characterized by an enzymatic core and by several ancillary subunits that increase biochemical heterogeneity and determine specific biological functions (Chan and Morey, 2019; Margueron and Reinberg, 2011; Pasini and Di Croce, 2016). The PRC1 core is formed by the E3 ligases RING1A or RING1B that, by interacting with the products of one of the six *Pcgf* paralogue genes (PCGF1-6), catalyse the mono-ubiquitination of Histone H2A at Lysine 119 (H2AK119ub1) (Blackledge et al., 2014; Gao et al., 2012; Wang et al., 2004). The PRC2 core is composed by two mutually exclusive methyltransferases EZH1 and EZH2 that, by associating to the scaffold proteins SUZ12 and EED, catalyse mono-, di- and tri-methylation of Histone H3 Lysine 27 (H3K27me1, me2 and me3) (Ferrari et al., 2014; Lavarone et al., 2019; Margueron et al., 2008; Shen et al., 2008). Both H2AK119ub1 and H3K27me3 are specifically enriched at repressed CpGi containing promoters and their loss correlates with increased transcriptional activity of target genes. The absence of either PRC1 or PRC2 activity results in developmental failure at pre- and post-implantation stages respectively (Faust et al., 1998; O’Carroll et al., 2001; Pasini et al., 2007; Posfai et al., 2012). In contrast, PRC1 loss of function in adult tissue severely compromises homeostasis which is not phenocopied by loss of PRC2 (Chiacchiera and Pasini, 2017).

The presence of several ancillary subunits determines the existence of many different PRC1 and PRC2 sub-complexes that may confer specific molecular properties and biological functions. PRC2 exists in two major forms: PRC2.1 and PRC2.2. PRC2.1 is characterized by the presence of Polycomb Like subunits (PHF1, MTF2 and PHF19) which confer affinity of the complex to recognise unmethylated CpG islands, and either EPOP or PALI1 (Beringer et al., 2016; Conway et al., 2018). PRC2.2 is characterized by the AEBP2 and JARID2 subunits, where JARID2 provides affinity to PRC2.2 to bind directly to H2AK119ub1 (Blackledge et al., 2014; Cooper et al., 2016; Kalb et al., 2014). PRC1 can instead exist in six distinct complexes (PRC1.1-6) characterized by six mutually exclusive PCGF paralogue subunits (PCGF1-6) (Gao et al., 2012; Hauri et al., 2016). PRC1.2 and PRC1.4 complexes are defined as canonical PRC1 (cPRC1) by the presence of CBX subunits that can bind H3K27me3, implying cPRC1 dependency on PRC2 activity (Blackledge et al., 2014; Cao et al., 2002; Tavares et al., 2012). PRC1.1, PRC1.3, PRC1.5 and the PRC1.6 forms exclude CBX proteins by associating with RYBP (or its paralogue YAF2), do not recognize H3K27me3 and their activity is independent of PRC2. These PRC1 forms are defined as variant PRC1 (vPRC1) and are tethered to target loci by intrinsic DNA binding activities. This includes PRC1.1 recognition of unmethylated CpG di-nucleotides by the KDM2B subunit (Farcas et al., 2012); PRC1.6 recognition of E-BOX and E2F DNA elements by the MAX/MGA and E2F6/DP dimers stably associated with the complex (Huang et al., 2018; Scelfo et al., 2019; Stielow et al., 2018); PRC1.3 (and likely PRC1.5) by the recognition of an E-BOX variant directly bound by the USF1/2 transcription factors that can interact with and recruit the PRC1.3 complex to chromatin (Scelfo et al., 2019). Overall, this involves the cooperative activity of both cPRC1 and vPRC1 forms at repressed sites together with the exclusive presence of vPRC1 forms (PRC1.6 and PRC1.3) at several highly expressed genes. While PcG repressed loci display abundant H2AK119ub1 decoration, active vPRC1 targets are characterized by a low-to-absent H2AK119ub1 deposition (Scelfo et al., 2019).

Although the role of vPRC1 complexes in transcriptional regulation remains unclear, these observations suggest that H2AK119ub1 should play a major role in establishing transcriptional repression. Such a model implies an initial deposition of H2AK119ub1 that enhances PRC2 stability, H3K27me3 deposition, recruitment of cPRC1 and establishment of PcG repressive domains (Blackledge et al., 2015). However, the central role of H2AK119ub1 in establishing PcG mediated repression remains controversial. While different reports provided evidences that H2AK119ub1 is required for the repression of PcG targets without altering PcG mediated chromatin higher order structures (Kundu et al., 2017), others have shown that the lack of H2AK119ub1 deposition is dispensable for homeotic gene regulation during *Drosophila melanogaster* development (Pengelly et al., 2015). Similarly, mice bearing an inactive RING1B point mutation (I53A) delayed the embryonic lethality of *Ring1B* KO mice from E10.5 to E15.5 (Illingworth et al., 2015).

In human tumours H2AK119ub1 deposition is enhanced by frequent inactivating mutations of the H2AK119ub1 specific deubiquitinase BAP1 (Carbone et al., 2013). Therefore, defining the central role of H2AK119ub1 in mediating transcriptional repression, and its relationship with PRC2 activity, remains an essential question to be addressed. Here we have developed an inducible system in mouse Embryonic stem cells (ESC) that allows us to dissect the contribution of H2AK119ub1 in regulating PRC1 and PRC2 mediated repression. Using a RING1B I53S catalytically inactive mutant, we showed that lack of H2AK119ub1 deposition, in the absence of cPRC1 and vPRC1 biochemical disruption, massively induced the transcriptional activation of PcG repressed targets with minimal indirect effects. Mechanistically this implied a strong destabilization of PRC2 complex activity that resulted in compromised H3K27me3 deposition that preferentially involved the H2AK119ub1-dependent PRC2.2 form. Finally, reduced H3K27me3 activity induced complete cPRC1 displacement from chromatin with minor effects on vPRC1 recruitment. Overall, these results place H2AK119ub1 deposition as the central modification for PcG-mediated control of transcriptional repression.

## Results

### Expression of RING1B I53S Missense Mutation Preserves PRC1 Assembly but Results in Complete Loss of H2AK119ub1 *in vivo*

To unravel the contribution of RING1B catalytic activity in PRCs-mediated transcriptional repression, we took advantage of ROSA26:creERT2 RING1A^-/-^; RING1B^fl/fl^ conditional mouse ESCs (Endoh et al., 2008) and manipulated this line (from now on defined as parental) by integrating a vector that stably expressed a FLAG-HA-tagged version of wild type mouse RING1B (WT) or the RING1B missense mutations I53A and I53S (Figure 1A). Treatment of these ESC with 4-hydroxy Tamoxifen (OHT) will induce complete loss of endogenous RING1A/B protein levels in the parental line or leave the unique expression of exogenous RING1B WT or RING1B I53A/S in the engineered ESC clones (Figure 1A). Our previous analysis with this parental line identified 72 hours of OHT treatment as the earliest time point at which endogenous RING1B and H2AK119ub1 deposition were lost (Lavarone et al., 2019). Indeed, at this time point, endogenous RING1B levels were undetectable resulting solely in the expression of the exogenous counterparts (Figure 1B). Importantly, WT and mutant exogenous forms were expressed to the same levels of endogenous RING1B without affecting the expression levels of PRC1 components that define canonical (CBX7) and variant PRC1 forms (RYBP) (Figure 1B). While RING1B WT expression did not alter the overall H2AK119ub1 levels, expression of RING1B I53A and I53S mutants resulted in the global loss of H2AK119ub1 deposition at levels comparable with OHT treated parental cells (Figure 1B). While the I53A mutation has been previously shown to be hypomorphic with some residual activity of H2AK119ub1 deposition, the I53S was shown to be fully catalytic dead (Ben-Saadon et al., 2006; Buchwald et al., 2006; Elderkin et al., 2007; Illingworth et al., 2015; Tsuboi et al., 2018) as confirmed by the complete lack of H2AK119ub1 deposition observed in our model (Figure 1B). We therefore decided to perform all further molecular analysis with this mutant line. Co-immunoprecipitation experiments showed that RING1B I53S efficiently formed canonical and variant PRC1 forms demonstrating that lack of H2AK119ub1 deposition is not a consequence of PRC1 complexes disruption (Figure 1C). Genome-wide localization analyses for H2AK119ub1 extended these observations, demonstrating that RING1B I53S expression induced a complete loss H2AK119ub1 deposition at all PRC1 target loci (Figure 1D and Table S1). Importantly, the expression of RING1B WT perfectly maintained physiological levels of H2AK119ub1 to all target loci (Figure 1D-F). This result was further validated by real time quantitative PCR (RTqPCR) analyses at selected loci (Figure 1G). Overall, these results demonstrated that the expression of RING1B I53S preserved PRC1 complex formation but was completely catalytically dead *in vivo* at all PRC1 target loci.

**Figure 1.**
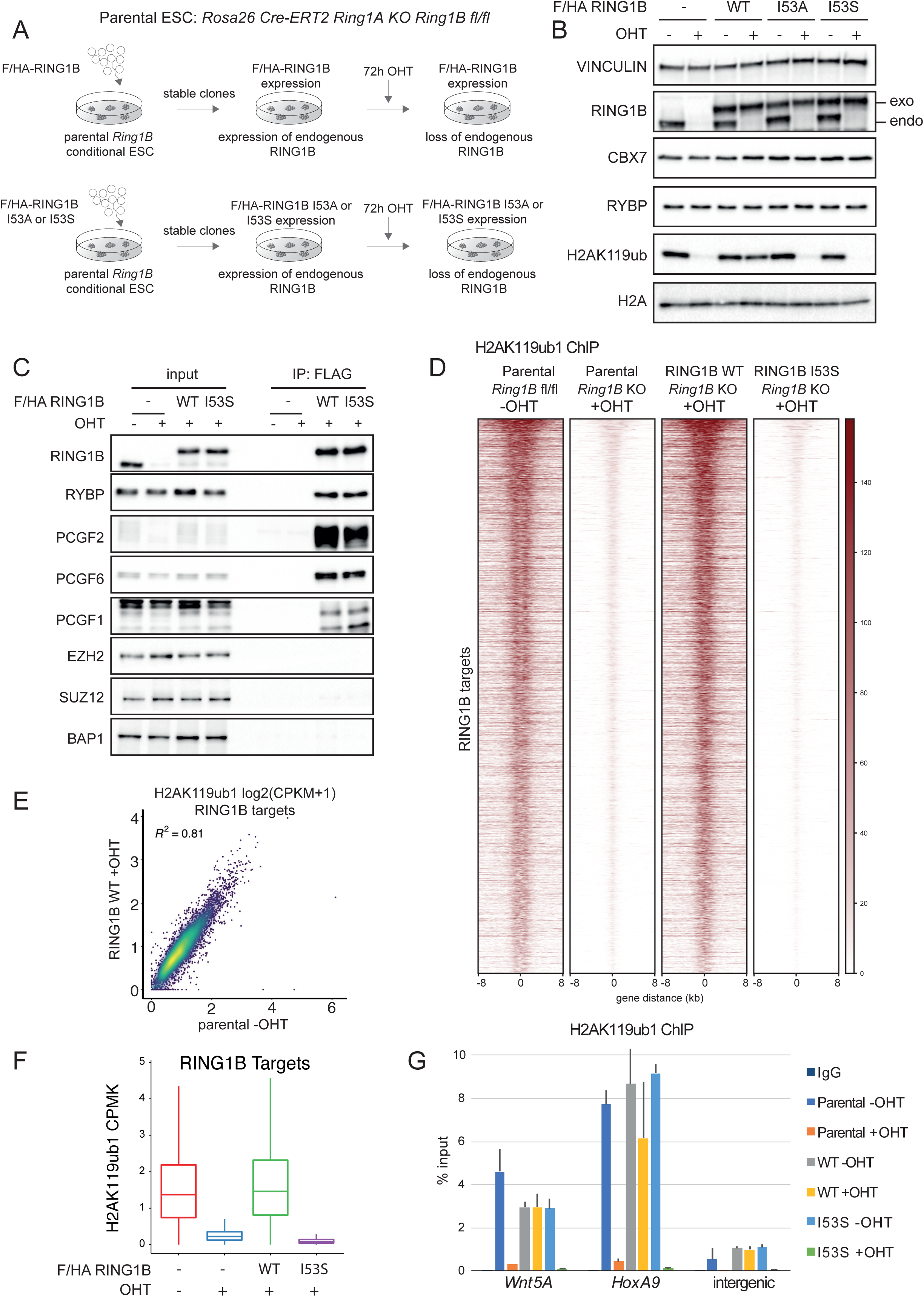
RING1B I53S is fully catalytically dead in vivo. A) Schematic representation of the strategy used for the generation of ROSA26:creERT2 RING1A^-/-^; RING1B^fl/fl^ conditional mESCs stably expressing FLAG-HA (F/HA)-tagged RING1B WT or I53A/S. B) Western blot analysis with the indicated antibodies of total protein extracts obtained from the specified cell lines upon 72 hours of treatment with OHT (+OHT) or EtOH (-OHT). Vinculin and Histone H2A were used as loading control. C) Co-immunoprecipitation analysis of nuclear extracts derived from FLAG-HA (F/HA) - tagged RING1B WT or I53S expressing cells upon 72 hours of treatment with OHT (+OHT) or EtOH (-OHT) using M2 affinity gel beads. FLAG-IPs in parental cells served as a negative control. D) Heatmaps representing normalized H2AK119ub1 ChIP-seq intensities ± 8Kb around the center of RING1B target loci in the indicated cell lines. E) Scatter plot showing the relationship between H2AK119ub1 CPMK levels (Counts Per Million per Kilobase) between parental EtOH treated (-OHT) and RING1B WT OHT treated (+OHT) cells in RING1B target loci. R^2^ represents the coefficient of determination of linear regression. F) Boxplots representing H2AK119ub1 ChIP-seq CPMK levels (Counts Per Million per Kilobase) in the indicated cell lines at RING1B target loci. G) ChIP RTqPCR analysis of H2AK119ub1 in the indicated cell lines at two specific Polycomb targets and one intergenic region. IgG served as a negative control.

### Deposition of H2AK119ub1 is Essential for PRC1-mediated Transcriptional Repression

Determining whether deposition of H2AK119ub1 is required for PRC1-mediated transcriptional repression remains an important open question. We took advantage of our model system to address this by performing RNA-seq analysis in parental, RING1B WT and RING1B I53S ESC at 72 hours after OHT treatment. Importantly, in the presence of endogenous RING1B expression (vehicle EtOH treatment: -OHT), ectopic expression of RING1B WT or I53S mutant did not alter the transcriptional landscape of ESC demonstrating that the I53S mutation does not exert any dominant negative effect on PRC1 activity (Figure 2A). Consistent with the repressive role of PRC1, addition of OHT to parental cells specifically induced the upregulation of a large number of repressed genes (Figure 2B and Table S2). While this effect is rescued by the ectopic expression of RING1B WT, the lack of H2AK119ub1 deposition - induced by RING1B I53S expression - perfectly phenocopied the transcriptional effects induced by global loss of PRC1 activity (Figure 2B and 2C). Few genes were also downregulated in both conditions suggesting common indirect effects (Figure 2D). Correlation plots further demonstrated high transcriptional concordance between parental and I53S ESC upon OHT treatment (Figure 2E), highlighting how genes enriched for H2AK119ub1 at their promoters are mostly upregulated (Figure 2E, red dots). Indeed, while the vast majority of upregulated genes are targets of PRC1 enzymatic activity, the few downregulated genes were mostly free of H2AK119ub1 deposition (Figure 2F). Overall these data directly place H2AK119ub1 deposition as an essential modification to maintain transcriptional repression of CpG-rich promoters.

**Figure 2.**
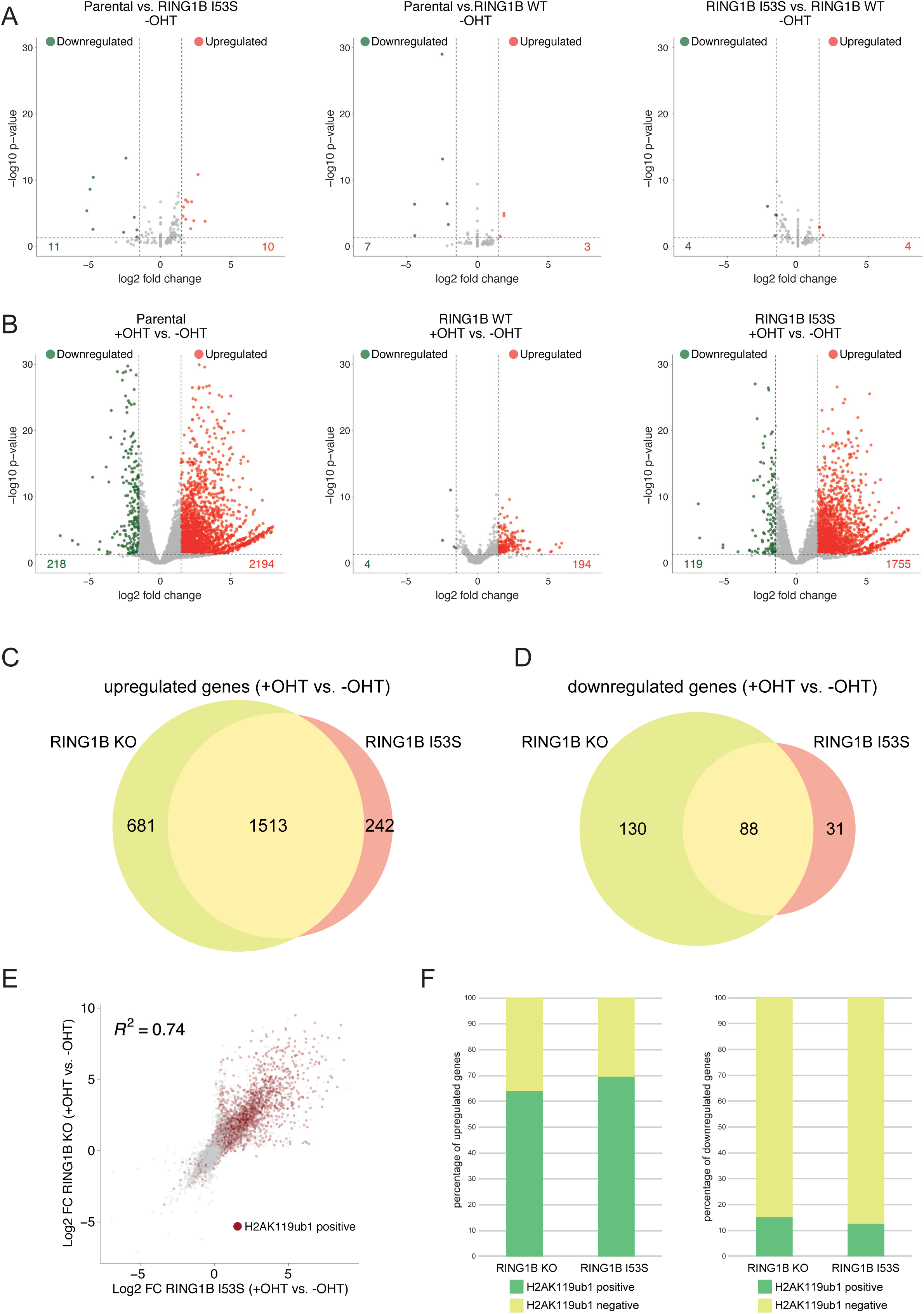
H2AK119ub1 is essential for PRCs-mediated transcriptional repression. A) Volcano plots of –log10 (P-value) against log2 fold change representing the differences in gene expression, related to RNA-seq analysis, in the indicated cell lines upon EtOH treatment (-OHT). Upregulated (Red) and Downregulated (Green) genes are highlighted. B) As in A upon OHT treatment (+OHT). C) Venn diagrams showing the overlap of upregulated genes between the indicated cell lines. D) As in C for downregulated genes. E) Scatter plot showing the relationship between log2 fold changes (FC) between the indicated cell lines at RING1B target loci. R^2^ represents the coefficient of determination of linear regression. Genes with promoters (± 2.5kb around TSS) containing H2AK119ub1 peaks are highlighted in red. F) Barplots showing the percentage of upregulated (Left) or downregulated (Right) genes with promoters (± 2.5kb around TSS) containing H2AK119ub1 peaks in the indicated cell lines.

### H2AK119ub1 Controls H3K27me3 Deposition by Regulating PRC2 recruitment

The mechanisms by which H2AK119ub1 could control transcriptional repression are still a matter of debate. H2AK119ub1 could be the end product of a PRC2-PRC1 functional crosstalk (canonical model) or the triggering modification that determines the recruitment of PRC1-PRC2 machineries. Using our system, we demonstrated that loss of H2AK119ub1 did not alter the expression levels of core PRC2 components (EZH2, SUZ12 and EED) as well as of sub-stoichiometric subunits that define the PRC2.1 (MTF2) and PRC2.2 (JARID2) variants (Figure 3A). We have noticed a modest reduction in H3K27me3 bulk levels in the absence of H2AK119ub1 deposition that suggest reduced PRC2 activity in agreement with previous reports (Blackledge et al., 2014; Fursova et al., 2019; Rose et al., 2016). Indeed, genome wide mapping of H3K27me3 deposition clearly showed a specific reduction of this modification demonstrating that PRC2 activity is severely affected by loss of H2AK119ub1 deposition (Figure 3B and 3C). This is consistent with the strong displacement of PRC2 from target loci observed by SUZ12 ChIP-seq analyses upon expression of RING1B I53S (+OHT) (Figure 3D and 3E). Overall, these results demonstrate that H2AK119ub1 is required to maintain efficient PRC2 recruitment and activity at its target sites.

**Figure 3.**
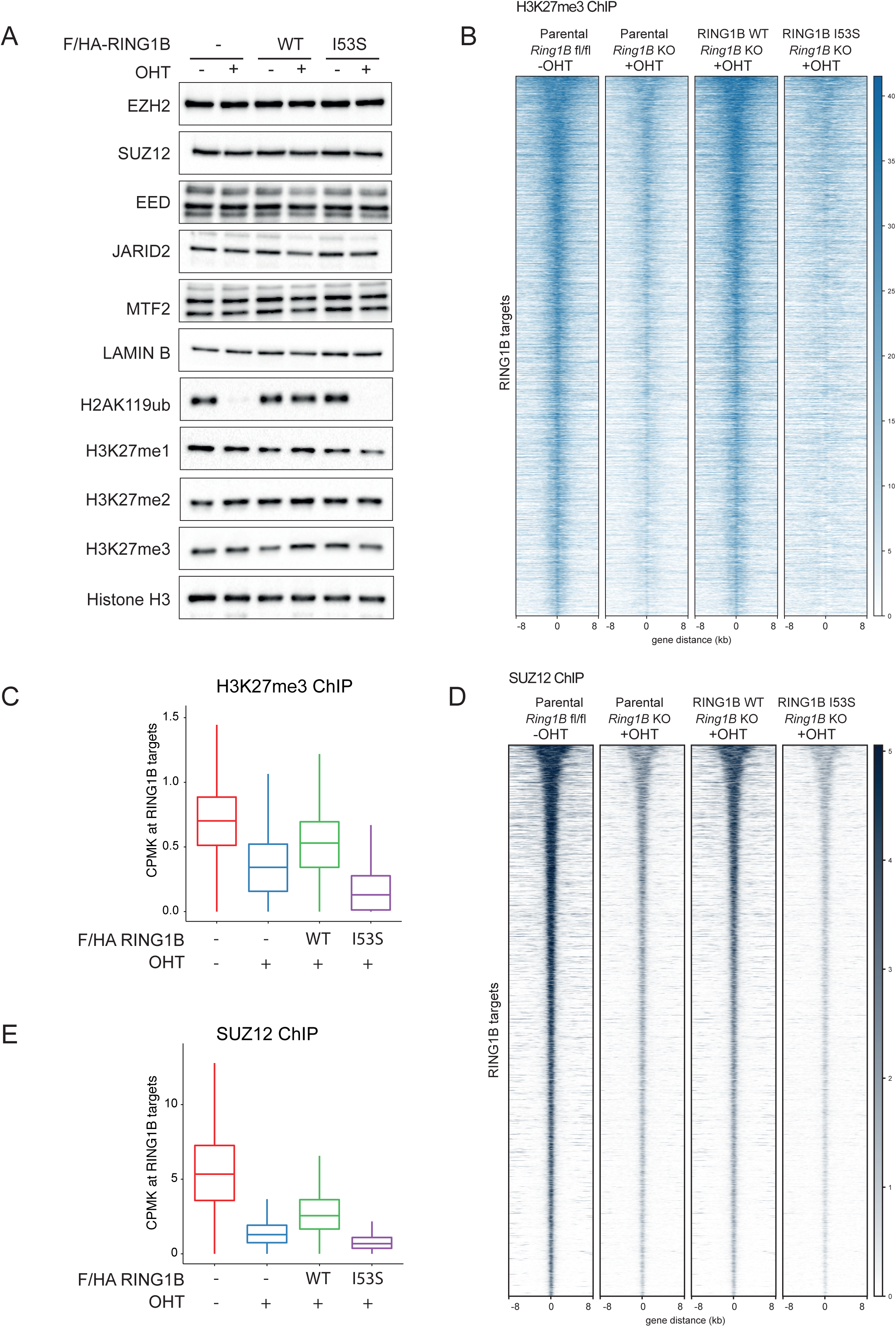
H2AK119ub1 deposition is required for PRC2 recruitment and activity. A) Western blot analysis with the indicated antibodies of protein extracts obtained from the specified cell lines upon 72 hours of treatment with OHT (+OHT) or EtOH (-OHT). LAMIN B and Histone H3 were used as loading controls. B) Heatmaps representing normalized H3K27me3 ChIP-seq intensities ± 8Kb around the center of RING1B target loci in the indicated cell lines. C) Boxplots representing H3K27me3 ChIP-seq CPMK levels (Counts Per Million per Kilobase) in the indicated cell lines at RING1B target loci. D) Heatmaps representing normalized SUZ12 ChIP-seq intensities ± 8Kb around the center of RING1B target loci in the indicated cell lines. E) Boxplots representing SUZ12 ChIP-seq CPMK levels (Counts Per Million per Kilobase) in the indicated cell lines at RING1B target loci.

### H2AK119ub1 Loss Triggers PRC2.1 and PRC2.2 Chromatin Displacement

The role that H2AK119ub1 plays in the recruitment of PRC2 also remains poorly characterized *in vivo*. Both H3K27me3 and H2AK119ub1 could serve as docking sites to stabilize PRC2 forms at target loci. Biochemical analyses had shown that JARID2 has direct affinity for H2AK119ub1 (Cooper et al., 2016) implying that PRC2.2 should be more dependent on this modification for its association at target promoters. At the same time, H3K27me3 can also serve as an affinity site for both PRC2.1 and PRC2.2 by EED recognition (Margueron et al., 2009). H3K27me3 binding by the WD40 repeats of EED stimulates PRC2 enzymatic activity allosterically inducing H3K27me3 spreading (Lee et al., 2018; Margueron et al., 2009). To shed light into this complex regulatory system, we tested whether loss of H2AK119ub1 deposition preferentially affected PRC2.1 and PRC2.2 target gene association by ChIP-seq analyses for MTF2 and JARID2, respectively. This revealed that both complex variants are affected by loss of H2AK119ub1 deposition (Figure 4A-D). However, differential intensity heatmaps and boxplot quantification highlighted a stronger displacement of JARID2 in contrast to MTF2 (Figure 4E and 4F) which is consistent with the affinity of JARID2 for H2AK119ub1. JARID2 and MTF2 displacement was further validated by locus specific RTqPCR analyses (Figure 4G and 4H), which also confirmed stronger MTF2 association at target loci in the absence of H2AK119ub1 deposition. Overall, these results demonstrate the central role of H2AK119ub1 in mounting both PRC2.1 and PRC2.2 at promoters, sustaining a positive feedback mechanism that allows the stabilization of PRC2 activities at target promoters.

**Figure 4.**
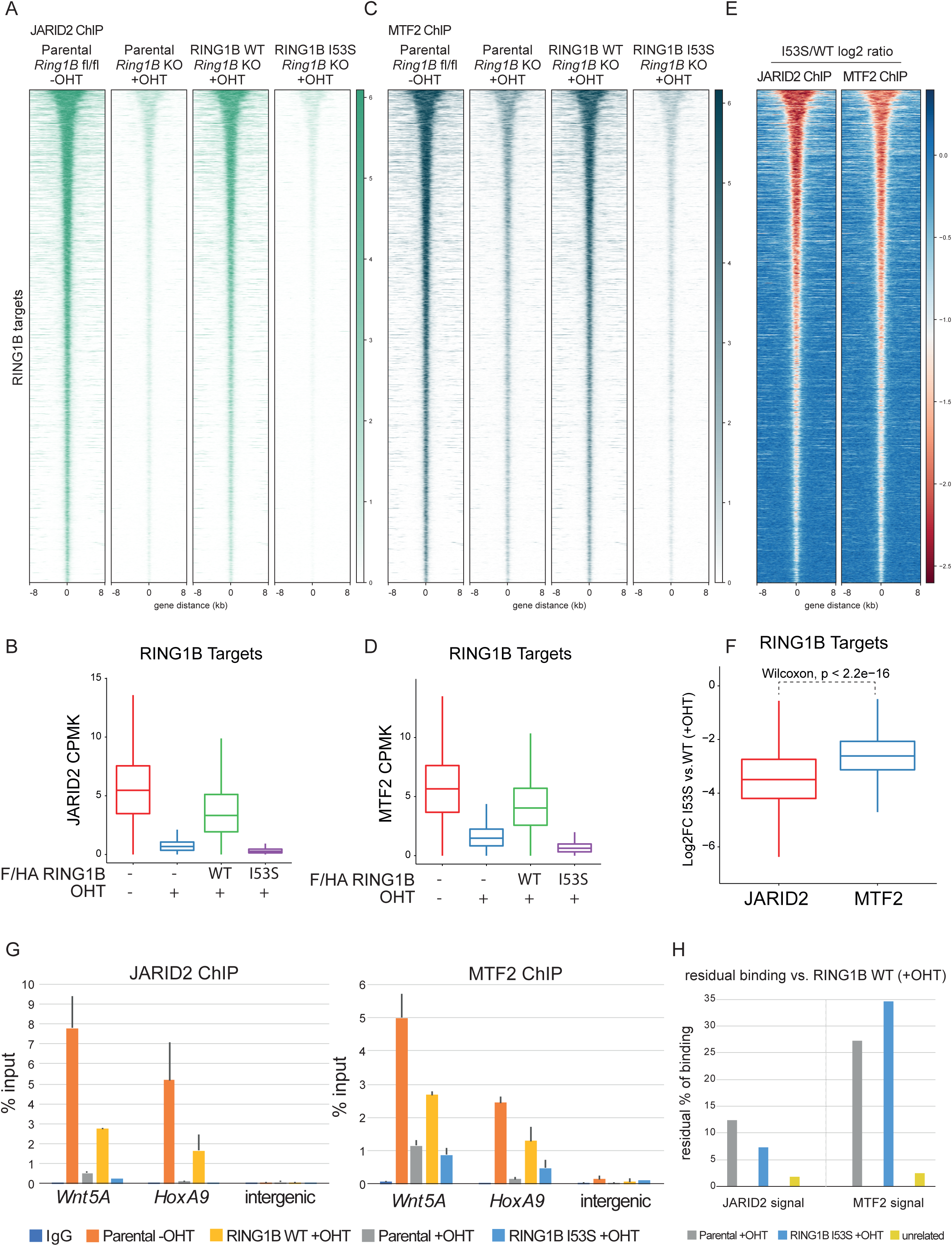
PRC2.1 and PRC2.2 variants sense H2AK119ub1 deposition differently. A) Heatmaps representing normalized JARID2 ChIP-seq intensities ± 8Kb around the center of RING1B target loci in the indicated cell lines. B) Boxplot representing JARID2 ChIP-seq CPMK levels (Counts Per Million per Kilobase) in the indicated cell lines at RING1B target loci. C) Heatmaps representing normalized MTF2 ChIP-seq intensities ± 8Kb around the center of RING1B target loci in the indicated cell lines. D) Boxplot representing MTF2 ChIP-seq CPMK levels (Counts Per Million per Kilobase) in the indicated cell lines at RING1B target loci. E) Heatmap representing the log2 ratio of JARID2 and MTF2 normalized ChIP-seq intensities at RING1B target loci between RING1B WT and I53S expressing cells. F) Boxplot representing the log2 ratio of JARID2 and MTF2 CPMK levels at RING1B target loci between RING1B WT and I53S expressing cells. G) ChIP RTqPCR analysis of JARID2 (Upper panel) and MTF2 (Bottom panel) in the indicated cell lines at two specific polycomb targets and one intergenic region. IgG served as a negative control. H) Histogram representing the residual percentage (%) of JARID2 and MTF2 binding in the RTqPCR analysis shown in D in the indicated cell lines.

### H2AK119ub1 Deposition Affects General RING1B Stability at Target Loci

The largest fraction of RING1B associated at target loci is dependent on H3K27me3 deposition. Only a minor residual fraction, corresponding to approximately 10% of RING1B signal, does not depend on this modification. However, this residual amount is highly active and is sufficient to preserve normal H2AK119ub1 deposition (Blackledge et al., 2014; Tavares et al., 2012). We therefore tested whether H2AK119ub1-dependent loss of H3K27me3 affected RING1B chromatin association. As expected, the vast majority of RING1B was displaced from target loci both in parental and RING1B I53S ESC treated with OHT (Figure 5A and 5B). This is not an intrinsic defect of the RING1B I53S mutant: RING1B I53S can assemble as a part of normal PRC1 complexes (Figure 1C) and can be efficiently recruited to target loci when H2AK119ub1 deposition is present (Figure 5C). Additionally, RING1B I53S fractionated in the nuclear insoluble fraction with the same efficiency of RING1B WT (Figure 5D). This demonstrated that neither localization nor chemical properties are altered by the mutation. This suggests that RING1B I53S displacement from target loci is a secondary effect of H2AK119ub1 and H3K27me3 loss. Indeed, RING1B was also globally displaced to a similar extent in EZH1/2 double KO ESC (Figure 5E and F), which lack H3K27me3 deposition but retain normal H2AK119ub1 levels (Lavarone et al., 2019). Overall, these results suggest that lack of H2AK119ub1 triggers a secondary displacement of the PRC1 complex from target loci that could depend on impaired PRC2 localization.

**Figure 5.**
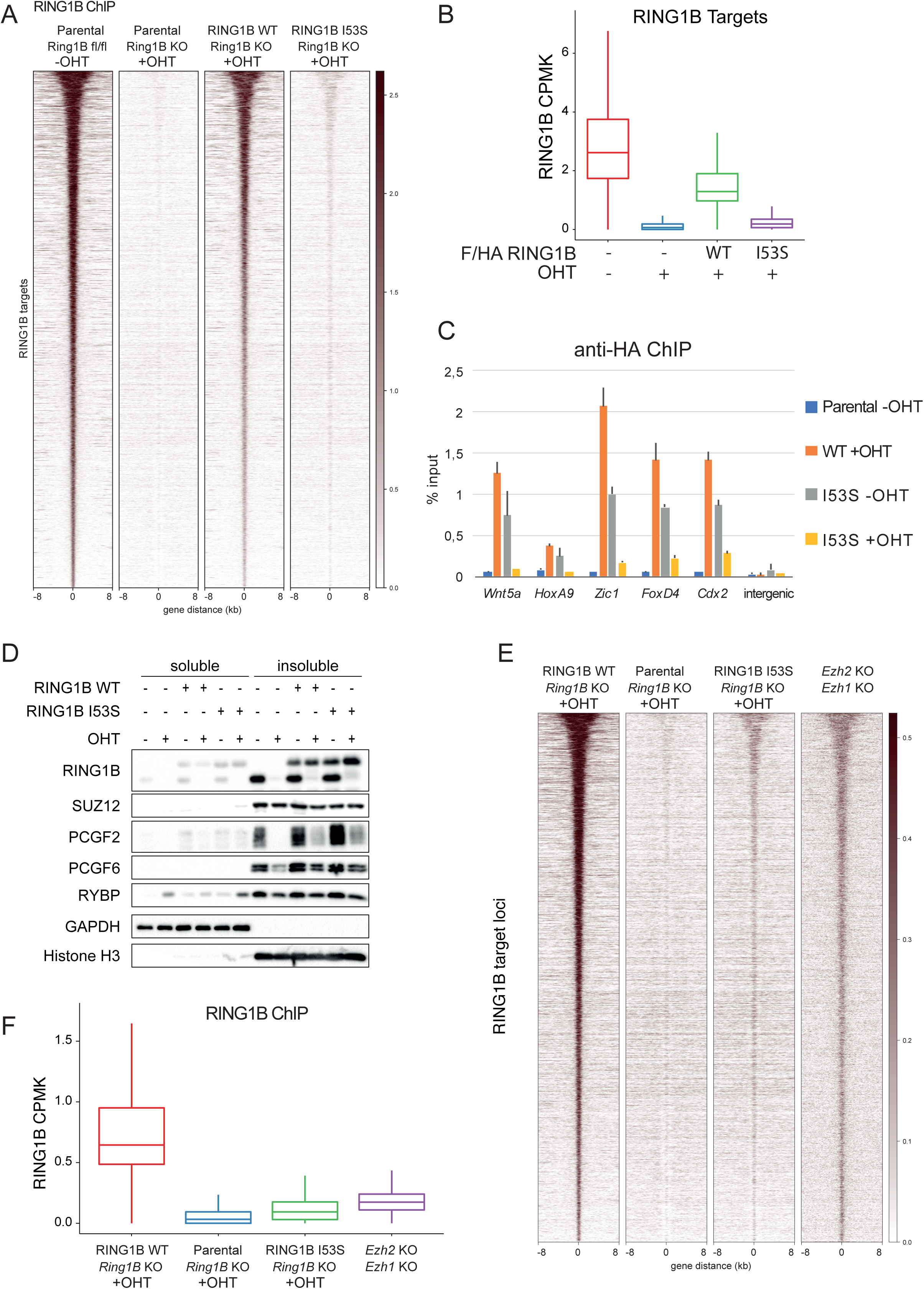
H2AK119ub1 affects RIN1GB chromatin stability. A) Heatmaps representing normalized RING1B ChIP-seq intensities ± 8Kb around the center of RING1B target loci in the indicated cell lines. B) Boxplots representing RING1B ChIP-seq CPMK levels (Counts Per Million per Kilobase) in the indicated cell lines at RING1B target loci. C) ChIP RTqPCR analysis of HA in the indicated cell lines at five specific polycomb targets and one intergenic region. Parental cells served as a negative control. D) Western blot analysis with the indicated antibodies of soluble and insoluble protein fractions obtained from the specified cell lines upon 72 hours of treatment with OHT (+OHT) or EtOH (-OHT). GAPDH and Histone H3 were used as positive controls for the soluble and insoluble fractions, respectively. E) Heatmaps representing normalized RING1B ChIP-seq intensities ± 8Kb around the center of RING1B target loci in the indicated cell lines. F) Boxplots representing RING1B ChIP-seq CPMK levels (Counts Per Million per Kilobase) in the indicated cell lines at RING1B target loci.

### H2AK119ub1 Loss Preferentially Affects cPRC1 vs. vPRC1

Chromatin association of cPRC1 has been extensively described to depend on H3K27me3 deposition (Scelfo et al., 2015). This affinity is conferred by the chromodomain of CBX proteins that are not present in vPRC1 forms (Bernstein et al., 2006; Cao et al., 2002). Indeed, RING1B residual binding in *Eed* KO ESC was shown to depend on vPRC1 forms compared to cPRC1 (Tavares et al., 2012). We therefore tested if cPRC1 binding is specifically affected in the absence of H2AK119ub1 deposition by analysing PCGF2 (cPRC1) and PCGF6 (vPRC1) behaviour. PCGF2 levels were destabilized in the absence of endogenous RING1B expression in agreement with previous reports (Scelfo et al., 2019) (Figure 6A). Importantly, RING1B WT and I53S rescued PCGF2 degradation to a similar extent, confirming that complex assembly - but not activity - is required for PCGF2 stability (Figure 6A). At the genome-wide level, PCGF2 association at target loci was preserved by the expression of RING1B WT but was strongly compromised by the absence of H2AK119ub1 deposition (Figure 6B and 6C). This is consistent with the reduced H3K27me3 levels observed in the absence of H2AK119ub1 deposition (Figure 3B-C). Unexpectedly, PCGF6 binding was also affected by the loss of H2AK119ub1, however to a much lower extent than PCGF2 (Figure 6B-D). These results were further confirmed by RTqPCR analyses showing strong loss of PCGF2 binding and proficient PCGF6 association at its target sites (Figure 6E). Importantly, we have previously shown that, while PCGF6 (PRC1.6) associates to a large set of target promoters together with PRC1.2 and PRC1.1, it also shares a substantial set of unique target sites with lower levels of H2AK119ub1 (Scelfo et al., 2019). To gain further insights related to the role that H2AK119ub1 plays in stabilizing PRC1.6 binding at target loci, we stratified PCGF6 targets based on the occupancy of PCGF proteins (Scelfo et al., 2019). Such analysis highlighted that, while PCGF6 binding was affected at sites that are co-occupied with PCGF2 and presented higher levels of H2AK119ub1 and H3K27me3 deposition, its binding at unique targets was not affected by loss of H2AK119ub1 deposition (Figure 7A-B). Similar to PRC2, vPRC1 also have an affinity for H2AK119ub1 (Kalb et al., 2014). Overall, our results demonstrate that while cPRC1 association is highly dependent on H2AK119ub1 deposition, vPRC1 target association can be stabilized by the high H2AK119ub1 levels found at repressed sites but retain intrinsic independent affinities for its target loci.

**Figure 6.**
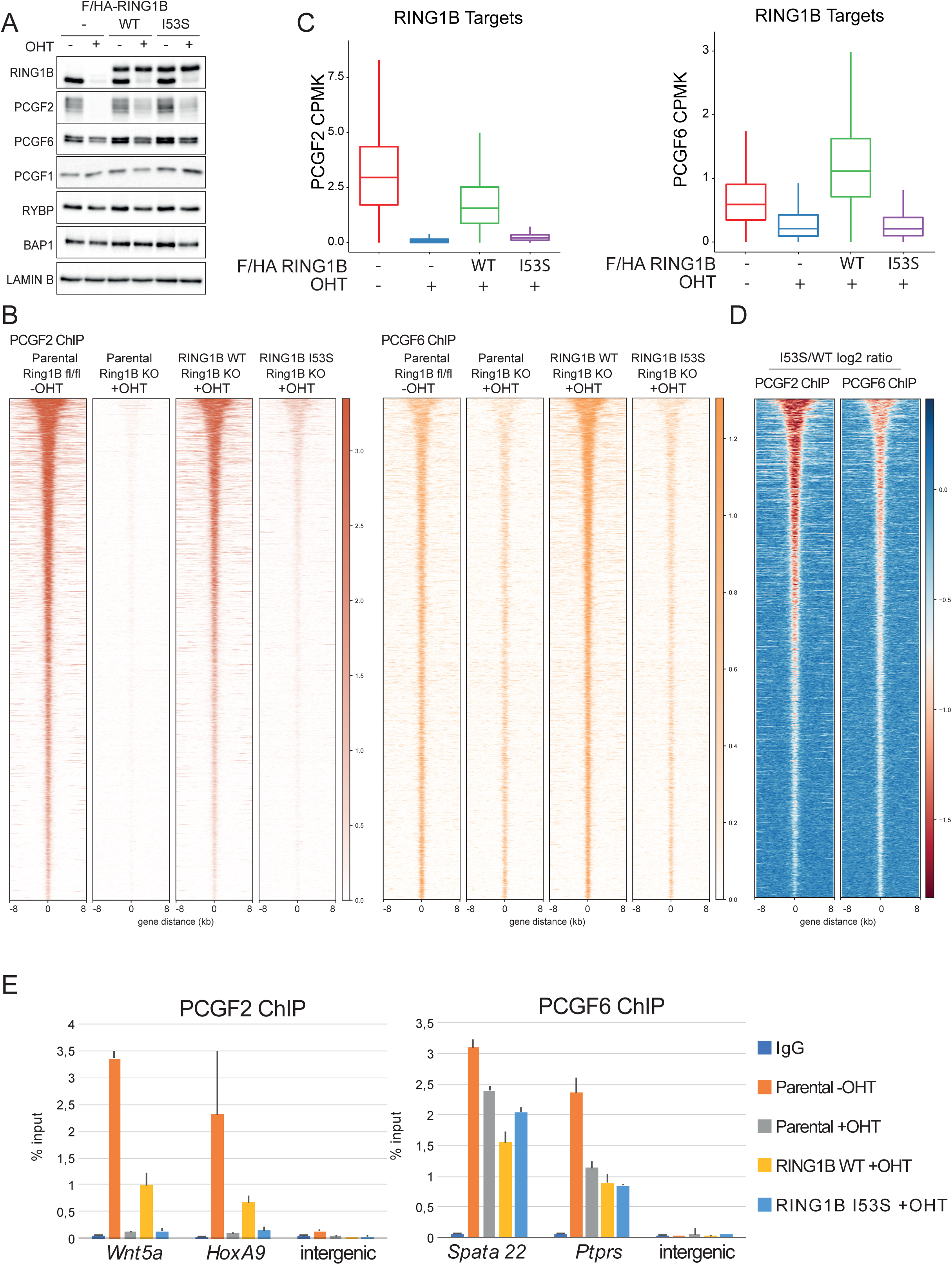
RING1B inactivation preferentially affects cPRC1. A) Western blot analysis with the indicated antibodies of nuclear protein extracts obtained from the specified cell lines upon 72 hours of treatment with OHT (+OHT) or EtOH (-OHT). LAMIN B served as a negative control. B) Heatmaps representing normalized PCGF2 and PCGF6 ChIP-seq intensities ± 8Kb around the center of RING1B target loci in the indicated cell lines. C) Boxplots representing PCGF2 ChIP-seq CPMK levels (Counts Per Million per Kilobase) (left panel) and PCGF6 (right panel) in the indicated cell lines at RING1B target loci. D) Heatmap representing the log2 ratio of PCGF2 and PCGF6 normalized ChIP-seq intensities at RING1B target loci between RING1B WT and I53S expressing cells. E) ChIP-qPCR analysis of PCGF2 (left panel) and PCGF6 (right panel) in the indicated cell lines at two specific PCGFs targets and one intergenic region. IgG served as negative control.

**Figure 7.**
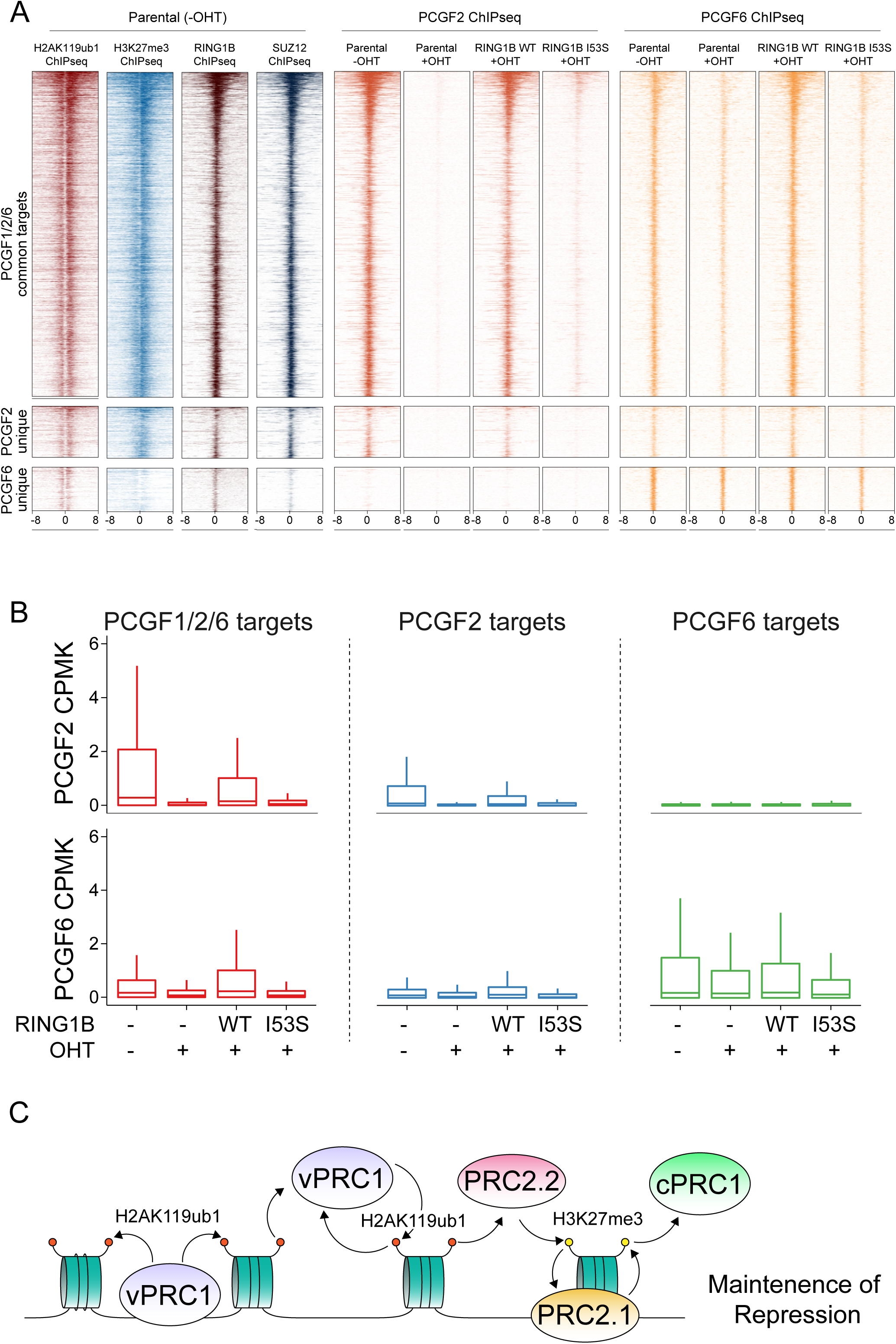
vPRC1 binding is more sensitive at sites of high H2AK119ub1 deposition. A) Heatmaps representing normalized H2AK119ub1, H3K27me3, RING1B, SUZ12, PCGF2 and PCGF6 ChIP-seq intensities ± 8Kb around the TSS of PCGF target genes in the indicated cell lines. B) Boxplots representing PCGF2 (upper panel) and PCGF6 (bottom panel) ChIP-seq CPMK levels (Counts Per Million per Kilobase) ± 250bp around TSS of PCGF target genes in the indicated cell lines. C) Model stressing the central role of H2AK119ub1 deposition in assembling both PRC2 and cPRC1 activities at repressed target genes but serving also as a positive feedforward mechanism to stabilize vPRC1 forms at the same sites.

## Discussion

The molecular mechanisms by which PRC1 and PRC2 activities control gene repression largely remain a matter of debate (Chan and Morey, 2019; Pasini and Di Croce, 2016). In this context, the role of cPRC1 vs. vPRC1 activity in mediating gene repression, as well as the direct role that H2AK119ub1 plays in this context, are an open important issue that requires further investigation. Here we have developed a simple model that generates inducible expression of a fully catalytically inactive form of RING1B (RING1B I53S in a RING1A null background) to dissect the contribution of H2AK119ub1 deposition to the structural properties of PRC1. This system allows monitoring of the acute effects induced by the loss of H2AK119ub1 deposition by preventing transcriptional adaptations and indirect effects that occur in constitutive PcG mutant ESC maintained in pluripotent conditions (Fursova et al., 2019; Scelfo et al., 2019). With this system, we showed that the expression of the catalytic inactive RING1B I53S neither affected the assembly of distinct PRC1 sub-complexes, nor its ability to associate with target promoters, but it did fail to maintain transcriptional repression of target genes to an identical extent as global RING1A/B deletion. This demonstrates the essential role of H2AK119ub1 deposition in maintaining transcriptional repression at genome wide levels in agreement with previous reports (Endoh et al., 2012). However, this differs from the role that was proposed for H2AK118ub1 (K119 in vertebrates) in *Drosophila melanogaster*. In this organism, H2AK118ub1 deposition was not required for correct spatio-temporal expression of homeotic genes during development (Pengelly et al., 2015) in contrast to the essential role played by H3K27me3 in the same context (Pengelly et al., 2013). Although PRC2 recognition of H2AK118ub1 is conserved in flies, these results suggest that loss of H2AK118ub1 is not sufficient to affect PRC2 activity at homeotic genes, perhaps due to redundant functions between PRC2.2 and PRC2.1, which is less affected by H2AK119ub1 loss. This is in line with the report that constitutive RING1B I53A mutant mice showed a substantial delay in embryonic lethality in contrast to *Ring1B* KOs (from E10.5 to E15.5) (Illingworth et al., 2015). Nonetheless, it remains unclear to what extent this developmental delay is directly caused by the lack of maintenance of target gene repression. Moreover, the hypomorphic properties of the RING1B I53A mutant (Tsuboi et al., 2018) used in this study, coupled to the expression of a proficient *Ring1A* allele, did not rule out the central role of H2AK119ub1 deposition in controlling transcriptional repression and early development. Now, our data has demonstrated the essential contribution of H2AK119ub1 deposition in PcG-mediated transcriptional repression, including HOX genes clusters. The different modalities of PcG complexes recruitment, together with the increased biochemical complexity of vPRC1 activity between flies and mammals, allows us to speculate that during vertebrate evolution a central role for H2AK119ub1 deposition in PcG-mediated transcriptional repression arose. The generation of more sophisticated and precise mouse models will become essential to dissect the contribution of H2AK119ub1-mediated control of transcriptional repression with respect to the more general role of PRC1 activities in development.

The work of several laboratories including ours have also highlighted that mammalian PRC1 exist in distinct biochemical forms (Gao et al., 2012) that are not exclusively associated with transcriptional repression (Fursova et al., 2019; Scelfo et al., 2019). Actively transcribed sites are specifically associated with PRC1.1, PRC1.3 or PRC1.6 forms. However, in this context, the role of H2AK119ub1 seems to be marginal as the vPRC1 forms associated with transcribed loci displayed low-to-absent H2AK119ub1 levels (Pivetti et al., 2019; Scelfo et al., 2019). The contribution of these PRC1 forms to active transcription remain poorly understood but is possible that these properties could have specific developmental roles that are not linked to H2AK119ub1 deposition and gene repression. Further studies in this direction are needed to dissect the repressive vs. the potential activatory role of specific vPRC1 forms. Our new data makes a large step in this direction demonstrating that, in the context of PcG-mediated transcriptional repression of CpGi containing promoters, H2AK119ub1 deposition serves as a central hub to maintain proper repression of target genes (summarized in Figure 7C model).

How PcGs establish and maintain transcriptional repression is also a matter of debate. Whether PRC2 controls PRC1 recruitment (canonical model) or PRC1 controls PRC2 (variant or non-canonical model) remains an open discussion (Chan and Morey, 2019; Pasini and Di Croce, 2016). Our data clearly showed that H2AK119ub1 function as a “glue” that keeps PRC1 and PRC2 machineries tethered to repressed loci at a genome-wide scale. Loss of H2AK119ub1 destabilizes PRC2 recruitment and H3K27me3 deposition, which induces displacement of cPRC1 forms, disrupting proper PcG repressive domains. The finding that this preferentially affected PRC2.2 activity is in agreement with the specific affinity of JARID2 for H2AK119ub1. Nonetheless, PRC2.1 was also affected in the absence of H2AK119ub1 suggesting a complex interplay between PRC2 variants. This could be an indirect consequence of reduced deposition of H3K27me3 levels together with a reduced affinity of PRC2.1 for target genes. Low H3K27me3 levels could reduce EED affinity for its target sites, compromising spreading and establishment of PcG domains (Lee et al., 2018; Oksuz et al., 2018). Reduced H3K27me3 deposition further correlated with specific cPRC1 displacement from all repressed target loci (represented by PCGF2 ChIP-seq), in agreement with cPRC1 being dependent on the deposition of this modification through the specific activity of CBX proteins (Bernstein et al., 2006; Cao et al., 2002). CBXs are excluded from vPRC1 forms by RYBP which were indeed less affected by lack of H2AK119ub1 deposition (exemplified by PCGF6 ChIP-seq). These results agree with the existence of distinct and specific mechanisms by which cPRC1 vs. vPRC1 forms are recruited to DNA and place vPRC1 activities upstream to cPRC1 (Figure 7C model).

Our data also showed that RING1B I53S became strongly destabilized from target genes in the absence of H2AK119ub1. This was primarily a consequence of the specific displacement of cPRC1 as a similar RING1B displacement was observed upon inactivation of PRC2 activity or upon expression of a catalytic-inactive PRC2 mutant that lack H3K27me3 deposition in the presence of normal H2AK119ub1 levels (Lavarone et al., 2019; Tavares et al., 2012). Indeed, significant binding of vPRC1 was still observed in the absence of H2AK119ub1. Importantly, in the presence of H2AK119ub1 deposition, RING1B I53S was recruited to target loci with the same efficiency as its *wild type* counterpart, demonstrating that its displacement was not an intrinsic DNA binding defect of this mutant. vPRC1 were also shown to retain *in vitro* affinity of H2AK119ub1 in ESC extracts (Kalb et al., 2014), suggesting a direct dependency on H2AK119ub1 deposition and a positive feedback loop mechanism that can further stabilize vPRC1 activity in addition to the PRC2-cPRC1 axis (summarized in Figure 7C model). Indeed, in the absence of H2AK119ub1 deposition, we observed a partial displacement of PCGF6, specifically at highly repressed sites. This observation further places direct H2AK119ub1 deposition in a central position to build up PcG repressive domains. This is consistent with the lack of transcriptional de-repression reported for the acute deletion of PRC2 activity in ESC (Riising et al., 2014) and further agrees with the lack of epistasis in early development and adult tissue homeostatic control between PRC1 and PRC2 activities (Chiacchiera et al., 2016a; Chiacchiera et al., 2016b).

Direct H2AK119ub1 deposition has also been linked to the development of several types of human cancers. This is specifically associated with the inactivation of the H2AK119ub1 specific de-ubiquitinase BAP1 that occurs with high frequency in Uveal Melanoma (∼45%) and Mesothelioma (∼22%) as well as in several other tumour types with lower frequencies like Atypical Spitz tumours (∼11%), Clear Cell Renal Cell Carcinoma (∼18%), Ovary (∼5%), and Colon-rectum (∼3%) (Carbone et al., 2013). BAP1 inactivation always resulted in strong accumulation of H2AK119ub1 levels (Campagne et al., 2019) and, based on the canonical model by which PRC2 controls upstream PRC1 activity, it was proposed that BAP1 deficient tumours could benefit from PRC2 inhibition (LaFave et al., 2015). Our data argues against this possibility placing H2AK119ub1 deposition in a central position to control PcG-mediated repression upstream to PRC2 and cPRC1 activities (Figure 7C model). This agrees with other reports that questioned the efficacy of PRC2 inhibition showing that viability of BAP1-deficient uveal melanoma cells is largely unaffected by PRC2 inhibition (Schoumacher et al., 2016). Based on our findings, we speculate that H2AK119ub1 deposition and vPRC1 activities could have a dominant role in controlling transcriptional repression also under pathological conditions with limited vulnerabilities within the PRC2-cPRC1 regulatory axis. Since we cannot exclude that BAP1 may also have additional roles that could be relevant in cancer progression (He et al., 2019), dissecting the contribution that vPRC1 forms play under pathological conditions of H2AK119ub1 deregulation will became a critical step for the future to define the tumour suppressive molecular properties of BAP1 and eventually uncover vulnerabilities for new strategies of intervention.

## Methods

### Cell lines and cell culture

ROSA26:creERT2 RING1A^-/-^; RING1B^fl/fl^ conditional mESC (Endoh et al., 2008) were engineered in order to stably express wild type or I53A/S RING1B. mESCs were grown on 0.1% gelatin-coated dishes in 2i/LIF-containing GMEM medium (Euroclone) supplemented with 20% fetal calf serum (Euroclone), 2 mM glutamine (Gibco), 100 U/ml penicillin, 0.1 mg/ml streptomycin (Gibco), 0.1 mM non-essential amino acids (Gibco), 1 mM sodium pyruvate (Gibco), 50 µM ß-mercaptoethanol phosphate buffered saline (PBS; Gibco), 1000 U/ml leukemia inhibitory factor (LIF; produced in-house), and GSK3β and MEK 1/2 inhibitors (ABCR GmbH) to a final concentration of 3 μM and 1 μM, respectively. Indicated cells were treated for 72 h with 0.5 μM 4-hydroxytamoxifen (OHT; or EtOH as vehicle) in order to delete *Ring1b* gene.

### Transfections

For stable clones generation, ROSA26:creERT2 RING1A^-/-^; RING1B^fl/fl^ conditional mESCs were transfected with pCAG vectors encoding 2xFlag-HA-tagged mouse wild-type or RING1B I53A/S using Lipofectamine 2000 (ThermoFisher Scientific), according to manufacturer’s instructions. Transfected cells were sub-cloned under puromycin selection (2 µg/ml) until the appearance of clones at day 10-12. Clones were screened by Western Blot and then selected for further analyses.

### Western Blot

For western blot analysis on total protein lysates, mESCs were lysed and sonicated in ice-cold S300 buffer (20 mM Tris-HCl pH 8.0, 300 mM NaCl, 10% glycerol, 0.2% NP40) and supplemented with protease inhibitors (Roche). Nuclear extracts were obtained as described below. Briefly, cells were resuspended in Hypotonic Buffer (10 mM TrisHCl pH 7.6, 10mM KCl, 1.5 mM MgCl2 and 0,340 M Sucrose supplemented with 2μg/mL Aprotinin, 1μg/mL Leupeptin,) for 15 minutes at 4° (C). Then 0.3% of 10% Triton X-100 was added to the solution and vortexed for 30 seconds followed by high speed centrifugation. The nuclear pellet was then washed with hypotonic buffer and solubilized in S300 buffer. Co-immunoprecipitations were performed on 2 mg nuclear extracts using M2 agarose beads (30 μl slurry for IP, A2220 Anti-FLAG M2 affinity gel) for 2 hours at 4°(C) Immunocomplexes were washed 5× with S300 buffer and eluted by competition with 1x Flag peptide (500 ng/ul; SIGMA) 2 times for 30 min at 16 °C and then resuspended in Laemmli sample buffer. Protein lysates were separated on SDS-PAGE gels and transferred to nitrocellulose membranes. After probing with the suitable primary and secondary antibodies, chemoluminescence signals were captured with the ChemiDoc Imaging System (Bio-Rad).

### Fractionation

Cellular pellets were lysed in 300μL pre-extraction buffer (20mM HEPES pH 7.5, 0.5% Triton X-100, 50mM NaCl, 3mM MgCl2, 300mM Sucrose, 2μg/mL Aprotinin, 1μg/mL Leupeptin, 1mM PMSF) and incubated at 4°C for 30 minutes. 150μL of suspension was removed and labeled “Total extract”. The remaining lysate suspension was clarified at 13,000RPM in a 4°C centrifuge for 10 minutes. Supernatant was transferred to a new tube and labeled “Soluble fraction”. The insoluble pellet was washed once in 1mL of pre-extraction buffer before resuspension in 150μL of pre-extraction buffer. All samples were boiled at 99°C for 5 min before sonicating 10 times (30 seconds on, 30 seconds off) at high intensity on a diagenode water bath sonicator.

### Antibodies

Western blot analyses were performed with: anti-Vinculin (V9131; Sigma-Aldrich), anti-BAP1 (D7W70; Cell Signaling), anti-Ezh2 (BD43 clone; homemade (Pasini et al., 2004)), anti-Suz12 (sc-46264; Santa Cruz Biotechnology), anti-Eed (AA19 clone; homemade (Bracken et al., 2003)), anti-flag (F3165; Sigma-Aldrich), anti-H3K27me1 (61015; Active Motif), anti-H3K27me2 (9728; Cell Signaling Technology), anti-H3K27me3 (9733; Cell Signaling Technology), anti-H2AK119ub (8240; Cell Signaling Technology), anti-H3 (ab1791; Abcam), anti-H2A (07-146; Sigma-Aldrich), anti-RING1B (homemade; (Chiacchiera et al., 2016a)), anti-RYBP (AB3637; Sigma-Aldrich), anti-CBX7 (ab21873, Abcam), anti-PCGF1 anti-PCGF2 and anti-PCGF6 (Pasini’s lab, homemade; (Scelfo et al., 2019)), anti-MTF2 (16208-1-AP, Proteintech), anti-JARID2 (13594, Cell Signaling Technology) and anti-LAMIN B (sc-6217, Santa Cruz Biotechnology).

ChIP assays were performed using: anti-Suz12 (3737; Cell Signaling Technology), anti-Ring1b (homemade; (Chiacchiera et al., 2016a)), anti-HA (12CA5 clone; homemade), anti-H3K27me3 (9733; Cell Signaling Technology), anti-H2AK119ub (8240; Cell Signaling Technology), anti-PCGF1, anti-PCGF2 and anti-PCGF6 (homemade (Scelfo et al., 2019), anti-MTF2 (16208-1-AP, Proteintech) and anti-JARID2 (13594, Cell Signaling Technology).

### Quantitative real-time PCR (qPCR)

Total RNA was extracted with the Quick-RNA™ MiniPrep extraction kit (Zymo Research) and retro-transcribed with ImProm-II™ Reverse Transcription System (Promega) according to the manufacturer’s instructions. Quantitative real-time PCR (qPCR) was carried out using GoTaq qPCR master mix (Promega) on CFX96 Real-Time PCR Detection System (Bio-Rad). Primer sequences are available upon request.

### Chromatin immunoprecipitation (ChIP)

ChIP experiments were performed according to standard protocols as described previously (Ferrari et al., 2014). For SUZ12, RING1B, PCGF1, PCGF6 and HA ChIPs, 1% formaldehyde cross-linked chromatin was sheared to 500–1000 bp fragments by sonication and incubated overnight in IP buffer (33 mM Tris-HCl pH 8, 100 mM NaCl, 5 mM EDTA, 0.2% NaN_3_, 0.33% SDS, 1.66% Triton X-100) at 4°C with the indicated antibodies (10 μg antibodies/ 500 μg chromatin). For histone modifications ChIPs, 250 μg of chromatin supplemented with 5% spike-in of S2 Drosophila chromatin (prepared in the same manner) and 5 μg of antibodies were used. The next day, chromatin lysates were incubated for 4 hours with protein-G sepharose beads (GE Healthcare). Beads were washed 3× with low-salt buffer (150 mM NaCl, 20 mM Tris-HCl pH 8, 2 mM EDTA, 0.1% SDS, 1 % Triton X-100) and 1× with high-salt buffer (500 mM NaCl, 20 mM Tris-HCl pH 8, 2 mM EDTA, 0.1% SDS, 1% Triton X-100), and then re-suspended in de-crosslinking solution (0.1 M NaHCO3, 1% SDS). DNA was purified with QIAquick PCR purification kit (Qiagen) according to manufacturer’s instructions. DNA libraries were prepared with 2–10 ng of DNA using an in-house protocol (Blecher-Gonen et al., 2013) by the IEO genomic facility and sequenced on an Illumina HiSeq 2000.

### RNA-seq

RNA-seq was performed following SMART-seq2 protocol (Picelli et al., 2014) with minor modifications. Briefly, poly-A containing mRNA molecules obtained from 1 μg of total RNA were copied into first-strand cDNA by reverse transcription and template-switching using oligo (dT) primers and an LNA-containing template-switching oligo (TSO). Resulting cDNA was pre-amplified with KAPA HotStart Taq enzyme (Kapa Biosystems) and then purified with Ampure beads (Agencourt AMPure XP-Beckman Coulter). Two nanograms of pre-amplified cDNA were tagmented with in-house produced Tn5 transposase and further amplified with KAPA HotStart Taq enzyme. After purification with Ampure beads, the quality of the obtained library was assessed by Bioanalyzer (High Sensitivity DNA kit, Agilent Technologies), prior to sequencing.

### ChIP-seq Analysis

Paired-end reads were aligned to the mouse reference genome mm10, or mm10 and dm6 for histone ChIP-Rx, using Bowtie v1.2.2 (Langmead et al., 2009) without allowing for multi-mapping (–m 1) and parameters -I 10 -X 1000. PCR duplicates were removed using samblaster (Faust and Hall, 2014). Ambiguous reads mapping to both mm10 and dm6 were discarded. Peaks were called using MACS2 v2.1.1 (Zhang et al., 2008) with parameters -f BAMPE --keep-dup all-m 10 30 -p 1e-10. A list containing the final RING1B and H2AK119Ub1 peaks used in the analyses, called in the WT +OHT cell line, can be found in Table S1. Genomic peak annotation was performed with the R package ChIPpeakAnno v3.15 (Zhu et al., 2010), considering the region +/– 2.5 kb around the center of the peak. PCGF target regions were obtain from (Scelfo et al., 2019) and liftOver to mm10. Peak lists were then transformed to gene target lists, and overlaps were performed using the R package VennDiagram v1.6.20 (Chen and Boutros, 2011).

For heatmap and intensity plot representation of ChIP-seq signal, BigWig files with input signal subtracted were generated using the function bamCompare from deepTools 3.1 (Ramirez et al., 2016) with parameters --ratio subtract –bs 50 --extendReads. To normalize for differences in sample library size, a scaling factor for each sample was calculated as (1/total mapped reads)*1000000 and was applied during BigWig file generation with the parameter --scaleFactors from bamCompare. For ChIP-Rx samples the scaling factor was calculated as described in (Orlando et al., 2014). For the spike-in samples, a second scaling factor was calculated based on the ratio mm10/dm6 reads of the input samples (one per cell line). The scaling factor from a particular input is applied to all its respective ChIP-Rx samples. This allows to correct any potential difference in the amount of spike-in added to the different pools of chromatin, which was one per cell line. Heatmaps were performed using the functions computeMatrix with settings reference-point --referencePoint center/TSS -b 8000 -a 8000 -bs 50, followed by plotHeatmap from deepTools excluding blacklisted regions by ENCODE (Consortium, 2012). For boxplot representation the function multiBigwigSummary (using BED-file and --outRawCounts options) from deeptools was used to calculate the average number of reads under peaks.

### RNA-seq Analysis

Reads were aligned to the mouse reference genome mm10 using STAR v2.7 without allowing multimapping reads (--outFilterMultimapNmax 1). PCR duplicates were removed using samblaster (Faust and Hall, 2014). Gene counts were calculated using featureCounts (Liao et al., 2014) with parameters -s 0 -t exon -g gene_name using Gencode M21 (GRCm38) annotation downloaded from (https://www.gencodegenes.org/mouse/). Differential expression analyses were performed using the R package DESeq2 v1.20 (Love et al., 2014) using default parameters. Log2FoldChanges and adjusted p-values were corrected using the apeglm (Zhu et al., 2019) and IHW (Ignatiadis et al., 2016) packages, respectively. Genes with an absolute log2 fold change of 1.5 and FDR < 0.05 were considered as differentially expressed (Table S2).

## Acknowledgments

We would like to acknowledge the IEO Genomic unit for support and all members of Pasini’s laboratory for helpful discussion. The work of the Pasini’s lab was supported by the Italian Association for Cancer Research, AIRC (IG-2017-20290) and by the European Research Council, ERC (EC-H2020-ERC-CoG-DissectPcG: 725268). DFP is a PhD student within the European School of Molecular Medicine (SEMM). DM was supported by a fellowship of FIRC and of European Institute of Oncology Foundation (FIEO). EC was supported by an iCARE2 fellowship of AIRC.

## Author contributions

S.T., E.L. and D.P. conceived the project. S.T and E.L. performed the experimental work. D.P.-F. performed the computational analysis. D.M, M.Z and E.C. provided support to the experimental work. S.T., E.L., and D.P. wrote the manuscript and edited the figures.

